# A single point mutation in the σ3 protein encoded by the reovirus S4 gene substantially attenuates the virulence of a highly myocarditic reassortant virus in neonatal mice

**DOI:** 10.64898/2026.02.25.707952

**Authors:** Meleana M. Hinchman, Andrew Miller, John S. L. Parker

**Affiliations:** Baker Institute for Animal Health, College of Veterinary Medicine, Cornell University, Ithaca, New York 14853; Department of Population Medicine and Diagnostic Sciences, College of Veterinary Medicine, Cornell University, Ithaca, New York 14853

## Abstract

Viral myocarditis is a major cause of heart damage, sudden death, and heart failure. Some strains of Mammalian Orthoreoviruses (REOV) cause myocarditis in newborn mice. This study examines the role of the REOV σ3 protein, encoded by the S4 gene, in modulating the virulence and myocarditic potential of a highly virulent T1L/T3DM2 reassortant virus that contains the M2 gene from the Type 3 Dearing (T3D) strain in the Type 1 Lang background.

We introduced single-point mutations in the double-stranded RNA-binding region of σ3 in the T1L/T3DM2 background. Our findings show that the K287T mutation in σ3 prevents the myocarditic phenotype and significantly reduces the virulence of T1L/T3DM2. Unlike the parental reassortant virus and a control reassortant mutant S4(R296T), infection of neonatal mice with the T1L-S4(K287T)/T3DM2 virus resulted in 100% survival, lower viral titers, particularly in the heart and spleen, and no gross or severe histological signs of myocarditis. This attenuation, despite similar *in vitro* growth and in vivo dissemination, indicates a tissue-specific replication deficit and highlights a key role for σ3 in the development of myocardial disease. The K287T mutation, unlike R296T, eliminates σ3’s capacity to inhibit protein kinase R (PKR) activation and to suppress NF-κB-driven transcription, leading to a strong innate immune response that likely controls viral replication and reduces cardiac damage. These results underscore the crucial role of the σ3 protein in modulating host innate immune responses and in influencing the outcome of REOV infection and myocarditis development.

**IMPORTANCE:** Viral myocarditis is a serious disease with life-threatening consequences, particularly in neonates and young animals. Infection of neonatal mice with certain strains of mammalian reovirus induces myocarditis. A reassortant virus containing the M2 gene from the Type 3 Dearing Strain in an otherwise Type 1 Lang background causes severe myocarditis in 100% of neonatal mice. Here, we show that a single point mutation in the S4 gene, which abolishes the capacity of the encoded σ3 protein to inhibit protein kinase R (PKR) and to suppress transcription of NF-κB-dependent genes, significantly attenuates the lethality of the T1L/T3DM2 reassortant and significantly reduces the severity of myocarditis. These findings highlight the importance of viral innate immune suppression in the induction of myocarditis.

## INTRODUCTION

Viral infection is a leading cause of myocarditis, characterized by inflammatory cell infiltration into the myocardium, loss of myocardial cells, calcium deposits, arrhythmias, and, in some cases, heart failure. The resulting damage can cause sudden death in children and young adults, especially males (1). Several viral infections are linked to myocarditis, including Coxsackievirus, Adenovirus, Influenza virus, Parvovirus, and COVID-19 virus (1, 2). Early diagnosis of viral myocarditis remains difficult because clinical signs, especially in infants and young children, can be nonspecific (3). Confirming the diagnosis requires an endomyocardial biopsy, an invasive procedure. Viruses can directly damage myocardial cells, leading to inflammation and myocarditis (4, 5). However, an excessive inflammatory response can also cause damage. Although many viral agents can cause myocarditis, the disease occurs sporadically, and the viral and host factors that determine whether a viral infection results in myocarditic damage are not well understood.

Mammalian Orthoreoviruses (REOV) are non-enveloped viruses with a genome of 10 segments of double-stranded RNA (dsRNA). Some REOV strains cause myocarditis in neonatal mice (6). Several viral genes, including M1, M2, and S4, have been linked to REOV-induced myocarditis (7–9). The T1L strain causes myocarditis in approximately 30-50% of infected mice, and infection is usually not fatal. Recently, Dina Zita and colleagues reported that a reassortant virus carrying the M2 gene from the Type 3 Dearing (T3D) strain in the T1L background caused severe myocarditis after oral inoculation of newborn mice (10). The T1L/T3DM2 reassortant was also significantly more virulent, with 90% lethality by day 7, compared with 100% survival in T1L-infected animals (10). These findings led the authors to conclude that the M2 gene plays a key role in REOV-induced myocarditis and that the T3D M2 allele imparts a lethal phenotype to the T1L REOV strain.

The M2 gene encodes the reovirus μ1 protein. μ1 assembles with the σ3 protein into heterohexamers that form a major component of the viral outer capsid (11, 12). As part of the outer capsid, μ1 undergoes conformational changes during virus entry, facilitating interaction with and penetration of the endosomal membrane, thereby allowing the viral core particle to enter the cytoplasm (11, 13, 14). μ1 also modulates NF-κB and IRF3 signaling (15, 16), limits RIP3-dependent necroptosis (17), and plays a role in REOV-induced apoptosis (18). In addition to protecting μ1 from proteolysis during virus entry, σ3 is a double-stranded RNA-binding protein that prevents activation of protein kinase R (PKR) and inhibits NF-κB signaling (11, 19). By mutating residues on σ3’s surface predicted to interact with dsRNA, we identified several residues that abolish dsRNA binding yet surprisingly do not block σ3’s capacity to inhibit PKR, indicating that dsRNA-binding is not necessary for PKR inhibition (20). Mutation of K287T within the dsRNA-binding region of σ3 abolished both dsRNA binding and σ3’s ability to prevent PKR activation. Viruses with this mutation grow to lower titers in tissue culture. We also found that, unlike WT T1L, T1L-S4(K287T) does not induce myocarditis despite replicating to similar titers in the heart (20). Because σ3 and μ1 interact in the cytoplasm, we wondered whether introducing the T3D M2 allele into the T1L background affects virulence and myocarditis partly through changes in σ3 function. To explore this hypothesis and the role of σ3 in modulating virulence and myocarditis, we generated σ3 mutants in the T1L/T3DM2 background.

Here, we show that a single point mutation in the reovirus σ3 protein eliminates lethality and reverses the severe myocarditic phenotype of the highly virulent T1L/T3DM2 reovirus reassortant.

## MATERIALS AND METHODS

### Cells and viruses

L929 cells were maintained in suspension in Joklik’s modified minimal essential medium supplemented with 5% fetal bovine serum, 2 mM glutamine, 1% antibiotic-antimycotic, and 0.05 mg mL-1 gentamicin. A549 cells were grown in Dulbecco’s modified Eagle medium (DMEM)/F-12 medium (Gibco) supplemented with 10% fetal bovine serum, 100 U mL^-1^ penicillin, 100 μg mL^-1^ streptomycin, and 1% L-glutamine. BHK-T7 cells were grown in DMEM supplemented with 10% fetal bovine serum, 1% L-glutamine, 100 U mL^-1^ penicillin, 100 μg mL^-1^ streptomycin, and 1 mg mL^-1^ G418 (Corning). Cells were incubated at 37°C with 5% CO2.

All viruses were recovered using reverse genetics (21, 22). Reovirus gene segments in pT7 vector plasmids were co-transfected with pCAG-FAST, pCAG-D1R, and pCAG-D12L into BHK-T7 cells using TransIT-LT1 (Mirus) according to the manufacturer’s instructions. The two S4 dsRNA mutants (K287T and R296T) were previously described (20). After transfection, cells were incubated at 37°C for 2 d, then frozen at -80°C. After two freeze-thaw cycles, lysates were cultured on L cell monolayers for plaque isolation. All viruses were purified, and sequences were verified by Sanger sequencing.

### Virus replication curves

L929 cells were infected with recombinant reoviruses at an MOI of 5 for single-cycle and at an MOI of 0.1 for multicycle growth curves. A549 cells were seeded and infected similarly at an MOI of 10 PFU per cell. At various time points post-infection, viral titers were measured by plaque assay on L929 cells.

### Virus purification

2 x 10^8^ L929 spinner cells were pelleted at 1000 rpm in the RC3B+ floor centrifuge and then suspended in 10 mL of virus at MOI 5 in complete Joklik’s media. After a one-hour incubation, the cells were resuspended to 400 mL and grown at 35°C until they reached approximately 65% CPE, as determined by trypan blue count on a hemocytometer. Cells were collected by centrifugation at 2300 rpm, and the pellet was resuspended in HO buffer (250 mM NaCl, 10 mM Tris, pH 7.4). Cell lysates were incubated with 1/100^th^ volume of 10% sodium deoxycholate for 30 minutes on ice, followed by sonication with Vertel XF and overnight centrifugation in a cesium chloride gradient from 1.25 to 1.45 g/cc in centrifuge tubes. Heavy virus bands were extracted from the gradient with a 21-gauge 1.5-inch needle and a 3 mL syringe. The virus sample was added to a presoaked 10K dialysis cassette (Thermo 66380) and dialyzed in 1x virion buffer (7.5 mM NaCl, 0.5 mM MgCl2, 0.5 mM Tris) with four exchanges: at 1-2 hours, 2 hours, 4 hours, and then an overnight to 2-day incubation with slow stirring at 4°C. The virus prep was then removed from the dialysis cassette, titrated, and sequence-confirmed along the S4 gene of T1L.

### Ethical approval for animal work

All animal work was conducted ethically in accordance with the US Public Health Service policy and approved by the Institutional Animal Care and Use Committee at Cornell University (IACUC no. 2019-0129). C57BL/6J mice were ordered from Jackson Laboratories and bred. Mice were housed in 11.5-inch × 7.5-inch IVC Polycarbonate Shoebox Cages for the duration of the experiment. Temperatures of 68–77 °F and humidity of 30-70% were maintained in the rodent room. Lights were turned on at 5:00 and off at 19:00 in the rodent room.

### Reovirus infections of neonatal C57BL/6J mice

Litters weighing 2-3 g per pup were orally gavaged with 50 μL of 10^7^ plaque-forming units (PFU) of reovirus T1L WT, T1L/T3DM2 reassortant, T1L-S4 (K287T)/T3DM2, T1L-S4 (R296T)/T3DM2, T1L-S4 (K287T), or T1L-S4 (R296T) in 1× PBS containing 10 μL green food color/mL (McCormick) using intramedic tubing (BD, 427401) via a 1 mL tuberculin slip-tip syringe (BD 309659) and a 30G 1/2 needle (BD 305106). Litters treated with 1× PBS containing 10 μL green food color/mL alone served as mock controls for the respective infection groups. Pups were weighed daily and identified with a skin marker (BrainTree Scientific KN-297). Neonates that lost weight for two consecutive days or appeared moribund were euthanized; otherwise, pups were culled at the end of the survival study and examined for gross signs of myocarditis. For titer, histology, cryo, and IF, pups were euthanized and tissues collected on days 1, 2, 4, and 7 post-inoculation.

### Tissue preparation for viral titration, cryopreservation, and paraffin embedding

Organs were collected from neonatal pups in ice-cold PBS. For viral titration, organs were placed into 2.0 mL tubes containing 3.0 mm zirconium beads (Benchmark D1032-30) and 1 mL of PBS. Organs were stored at -80°C until homogenized using the Benchmark Beadblaster 24 (model D2400). The homogenate was used for plaque assay on L cell monolayers in 6-well plates. Machine settings were 5K speed, 45 seconds, with 30-second rest intervals, repeated 4 times, then immediately refrozen.

### Cryo-sectioning and paraffin embedding

Hearts were first slowly perfused with PBS through a 27G 1/2 needle (BD 305109) attached to a syringe, then extracted for OCT compound (Tissue-TeK 4583) embedding or formalin fixation (Sigma HT501128) followed by paraffin embedding. Paraffin embedding and H&E staining were performed by the histology department of the Cornell University Diagnostics center. Other organs were harvested, placed in Petri dishes with ice-cold PBS, and then placed into cryomolds (Tissue-Tek 4566). Tissue was covered with OCT, frozen on the surface of a shallow 2-Methylbutane bath (Supelco MX0760) chilled over LN2 vapor, and then stored at - 80°C until ready for cutting. 10 µm cryosections were made using a microtome cryostat (Micron International GmbH HM505E) onto glass slides (Fisherbrand 1255015).

### Sample preparation for immunofluorescence

Ten-micron heart tissue sections were placed on microscope slides (Fisherbrand 1255015) and stored at -80°C. Slides were heated for 1 minute at 37°C to dry, then immediately immersed in pre-chilled methanol at -20°C for 30 minutes. After fixation, slides were rinsed in 1× TBS. Samples were permeabilized with 0.1% Triton X-100 in 1× PBS for 10 minutes, washed once in TBS for 5 minutes, and blocked for 1 hour at room temperature in blocking buffer (1% BSA and 10% normal donkey serum in PBS). Next, Frame-Seal chamber stickers (Bio-Rad SLF-0201) were placed around the tissue, and 30 µL of primary antibody, diluted in antibody solution (1% BSA in TBS), was added. Covers were carefully sealed over the chambers, and slides were incubated overnight at 4°C. The primary antibody was rabbit anti-reovirus VM1:VM6 polyclonal sera (1:30,000). After incubation, Frame-Seals were removed, samples were washed three times in TBS, and then incubated with secondary antibody labeled with fluorescent 488 at 1:500 (Jackson ImmunoResearch 711-545-152), diluted in blocking solution, for 1 hour in the dark at room temperature. Samples were washed three times in TBS for 5 minutes each. DAPI stain (Invitrogen R37606), diluted in PBS, was added to the tissues for 5 minutes in the dark, then washed twice with PBS for 5 minutes each. Samples were mounted with ProLong antifade mounting media (Invitrogen P36982) and cover glass (Globe Scientific INC 1404-10). Fourfold images were acquired on a Cytation 7 imager (Biotek Instrumentation, Inc.) using Gen5 software to analyze and compare cross-sections. DAPI-stained nuclei were imaged separately.

### Statistical analysis

All experiments were performed at least three times, and the data are presented as mean ± standard deviation (SD). Graphs and statistical analyses were performed using Prism^®^ GraphPad software, and the test type is described in the figure legends.

## RESULTS

### Viruses with mutations in the double-stranded RNA-binding region of σ3 reach titers similar to those of the parental T1L/T3DM2 reassortant

We previously demonstrated that the T1L-S4(K287T) recombinant virus was attenuated in growth and did not induce myocarditis in newborn mice (20). Therefore, we tested whether the K287T mutation in σ3 would attenuate the highly virulent and myocarditic T1L/T3DM2 reassortant (10). Using reverse genetics, we prepared the parental reassortant T1L/T3DM2, the T1L-S4(K287T)/T3DM2 mutant, and an additional control mutant, T1L-S4(R296T)/T3DM2. The R296T mutation in σ3 prevents dsRNA binding but does not affect its ability to inhibit PKR or NF-κB transcription (20, 23). The T1L-S4(K296T) virus has growth comparable to WT T1L and induces myocarditis (20). We first tested the growth capacity of the recombinant viruses in vitro. All viruses showed similar single-cycle growth kinetics in L929 cells (data not shown). Multiple growth cycles of the recombinant viruses in L929 and A549 cells were also similar (Fig. 1), indicating that, unlike our previous finding that the T1L-S4(K287T) virus was attenuated compared to T1L, the introduction of the S4(K287T) mutation into the T1L/T3DM2 background does not reduce in vitro growth.

**Figure 1.**
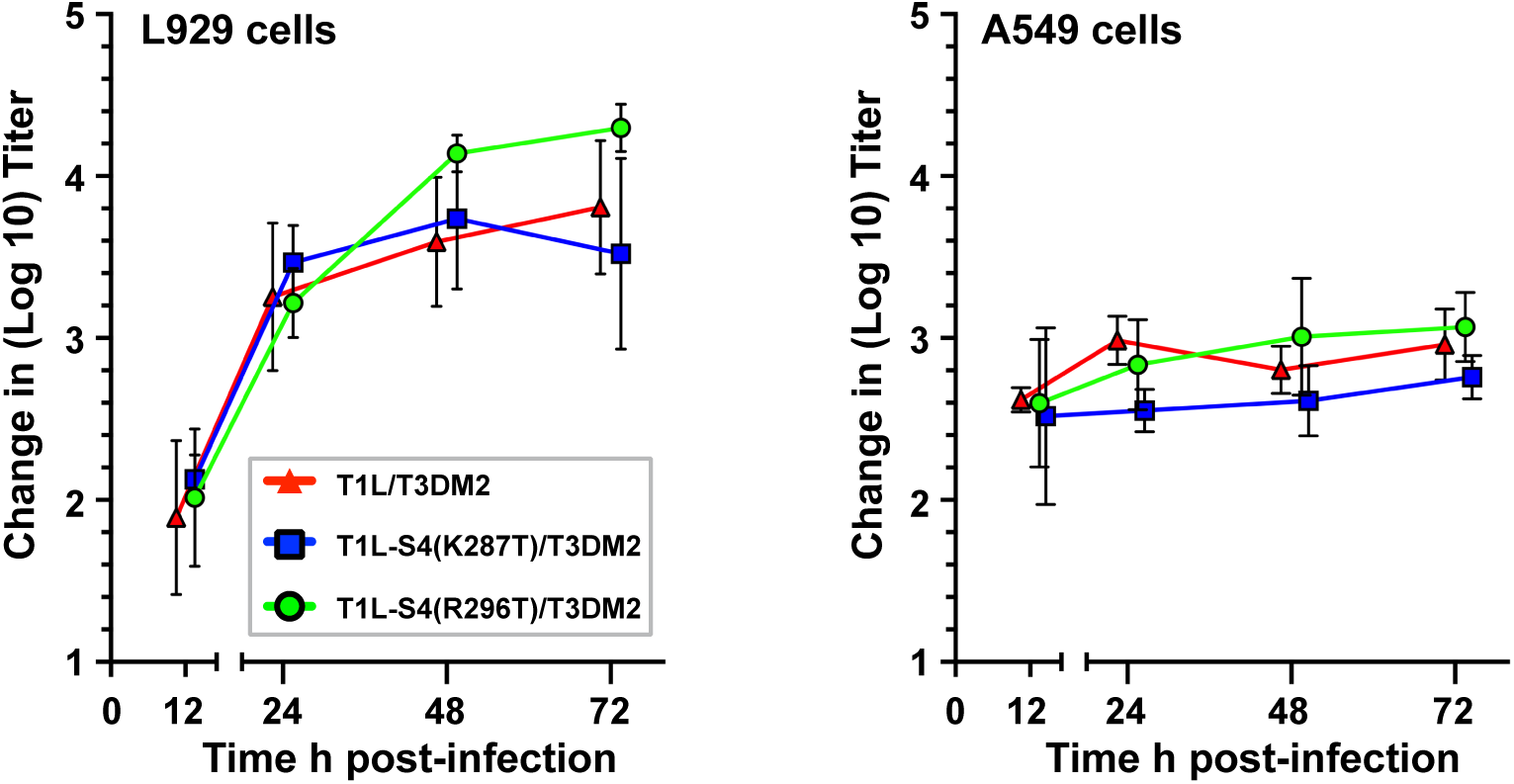
The TlL/T3DM2, T1L-S4(K287T)/T3DM2, and T1L-S4(R296T)/T3DM2 viruses exhibit similar growth kinetics in L929 and A549 cells. L929 and A549 cells were infected with purified, sequence-verified viruses at an MOI of 0.1 and 10, respectively. Viral titers were measured at the indicated time points. The mean and standard deviation of the change in log titer shown represent four independent biological experiments, each with two technical replicates.

The S4(K287T) mutation reduces the lethality of the T1L/T3DM2 reassortant. The T1L/T3DM2 reassortant was reported to be 90% lethal within 7 days after oral inoculation of newborn mice (10). Therefore, we compared the lethality of the T1L, T1L-S4(K287T), T1L-S4(R296T), T1L/T3DM2, T1L-S4(K287T)/T3DM2, and T1L-S4(R296T)/T3DM2 viruses. We found that, like T1L, oral inoculation of the T1L-S4(K287T) virus was non-lethal in C57BL/6 newborn mice (Fig. 2). Conversely, the T1L-S4(R296T) virus caused 33% mortality by day 20 post-inoculation, suggesting that the R296T mutation in σ3 increases the virulence of the T1L strain. Inoculation with the T1L/T3DM2 reassortant resulted in 50% lethality. A similar level of lethality was observed for the T1L-S4(R296T)/T3DM2 virus. However, all mice inoculated with the T1L-S4(K287T)/T3DM2 survived, indicating that the S4(K287T) mutation attenuates the virulence of the T1L/T3DM2 reassortant.

**Figure 2.**
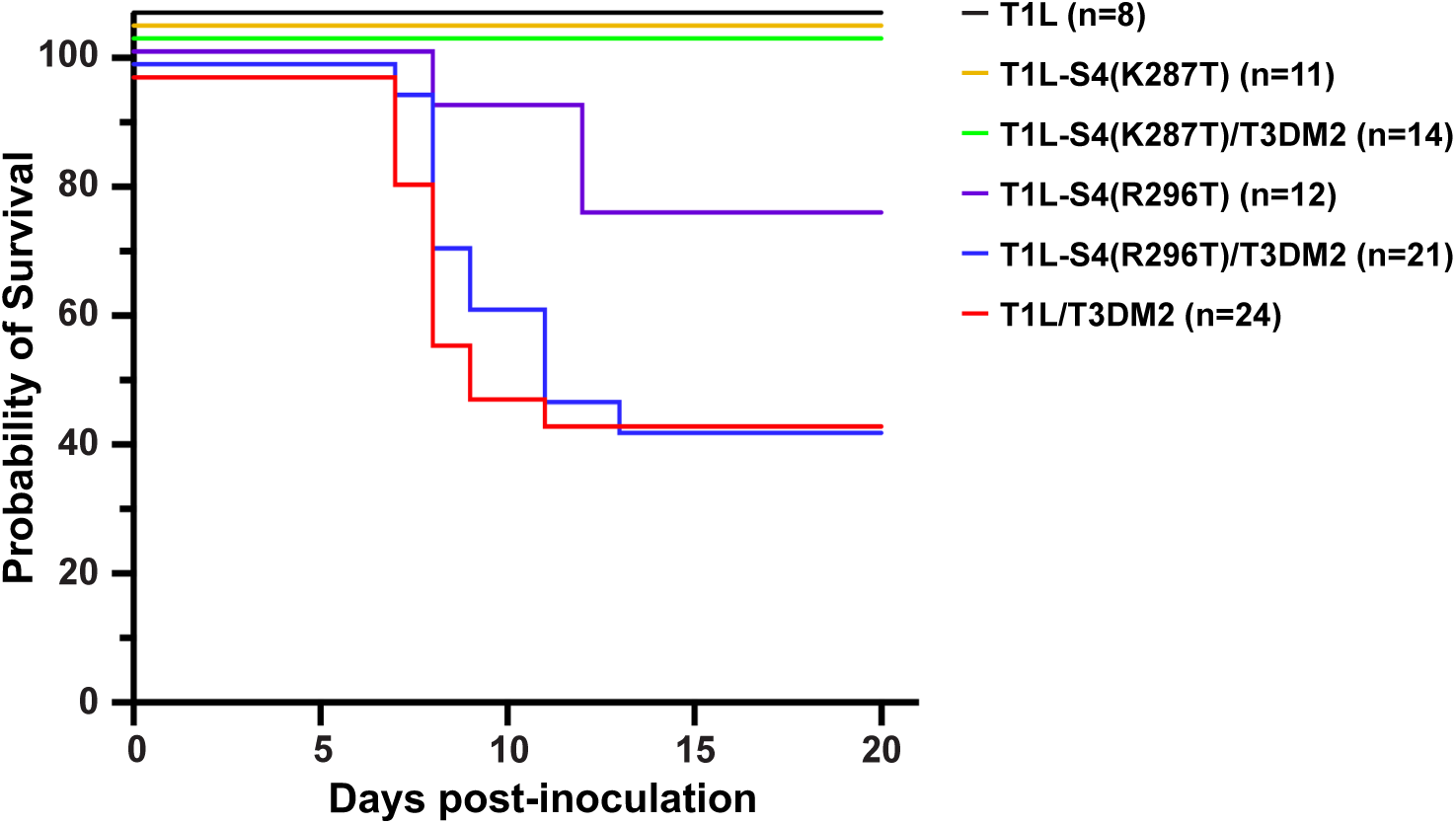
Introduction of the S4(K287T) mutation into the T1L/T3DM2 reassortant background significantly improves survival of neonatal mice after oral inoculation. Mouse pups were orally administered 1 × 10^7^ pfu of one of five REOV strains and were weighed daily for up to 20 days. The day of death or euthanasia was recorded. Pups that lost weight on two consecutive days were euthanized. All hearts were examined postmortem for gross myocarditic lesions. Significant differences between survival curves were assessed using the log-rank (Mantel-Cox) test. P-values were T1L vs T1L/T3DM2 = 0.0138, T1L vs T1L-S4(R296T)/T3DM2 = 0.0098, T1L/T3DM2 vs T1L-S4(K287T)/T3DM2 = 0.0011, and T1L-S4(R296T)/T3DM2 vs T1L-S4(K287T)/T3DM2 = 0.0007.

### The T1L-S4(K287T)/T3DM2 mutant spreads from the ileum but replicates at lower levels in the heart and spleen

Given differences in lethality, we measured viral loads in organs after oral inoculation with 10^7^ PFU of T1L/T3DM2, T1L-S4(K287T)/T3DM2, and T1L-S4(R296T)/T3DM2 (**Fig. 3**). In the ileum, peak viral titers occurred on day 1 and decreased on days 2, 4, and 7. However, on days 1 and 2, T1L-S4(K287T)/T3DM2 titers were significantly lower than those of the other two viruses. Despite lower viral loads in the ileum on days 1 and 2, the T1L-S4(K287T)/T3DM2 reassortant spread to secondary replication sites and showed titers comparable to those of the parental reassortant in the liver and brain on days 2 and 4. Unlike the parental reassortant and the T1L-S4(R296T)/T3DM2 mutant, the T1L-S4(K287T)/T3DM2 virus reached significantly lower peak titers in the heart and spleen by day 4, and these peaks were not maintained on day 7. Based on these findings, we conclude that the T1L-S4(K287T)/T3DM2 mutant exhibits tissue-specific replication limitations in the spleen and heart.

**Figure 3.**
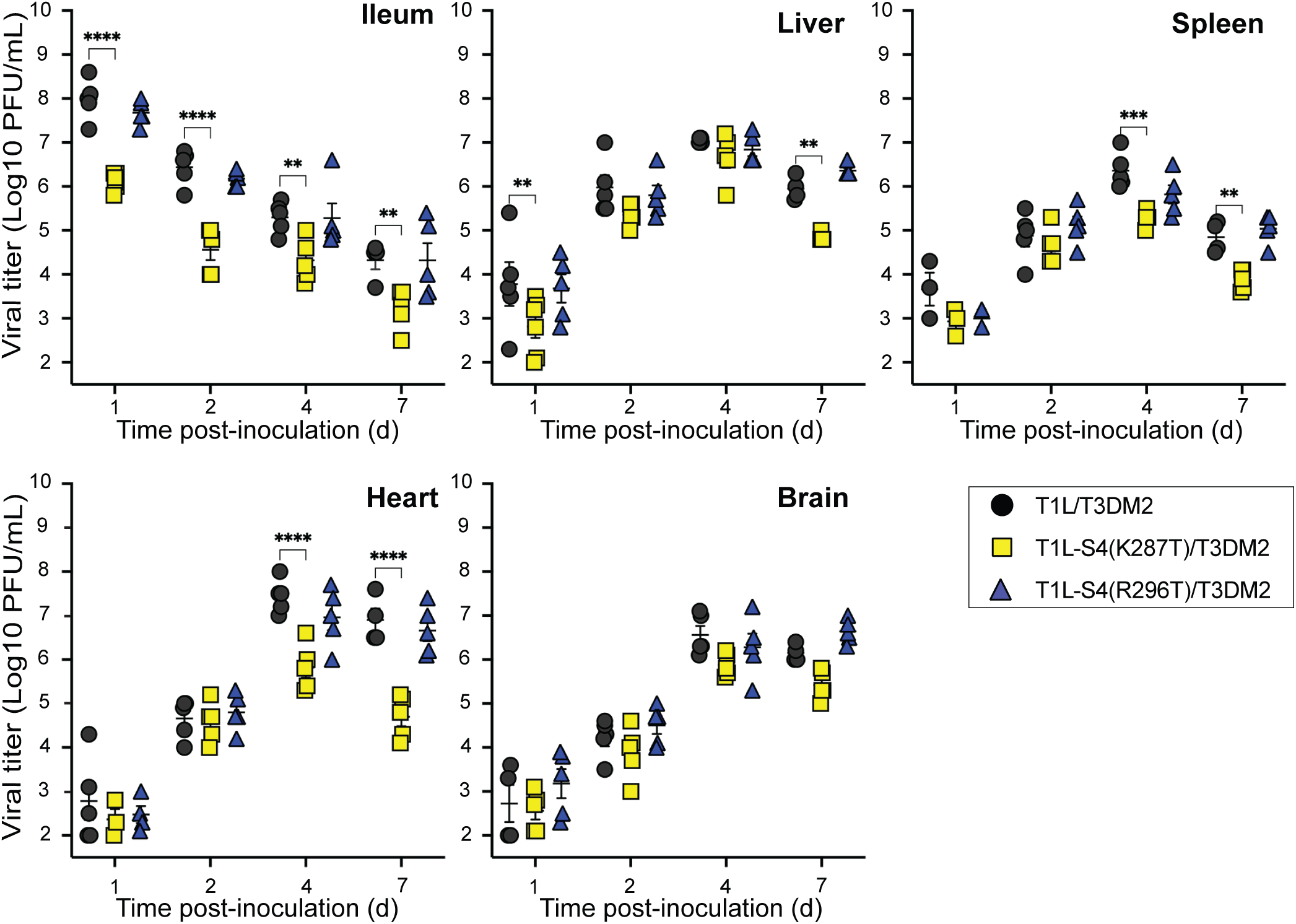
Compared with the T1L/T3DM2 parental reassortant, the T1L-S4(K287T)/T3DM2 virus shows lower viral titers in the intestine, spleen, and heart. Neonatal mice were inoculated orally with 10^7^ PFU of T1L/T3DM2, T1L-S4(K287T)/T3DM2, or T1L-S4(R296T)/T3DM2. At days 1, 2, 4, and 7, mice were euthanized, and the indicated tissues were collected. Viral titers in each tissue were determined by plaque assay (n = 3-5 per time point). A two-way ANOVA was used to compare the mean titers of the two mutant viruses with those of T1L/T3DM2 at each time point.

### The K287T mutation in σ3 reduces the ability of the T1L/T3DM2 reassortant to cause severe myocarditis in mice and lessens the severity of histological changes

By day 7 after oral inoculation, mice inoculated with the T1L/T3DM2 reassortant or T1L-S4(R296T) developed visibly abnormal hearts with large, striated lesions typical of myocarditis. In contrast, on day 21, mice inoculated with T1L-S4(K287T)/T3DM2 had visibly normal hearts (**Fig. 4**).

**Figure 4.**
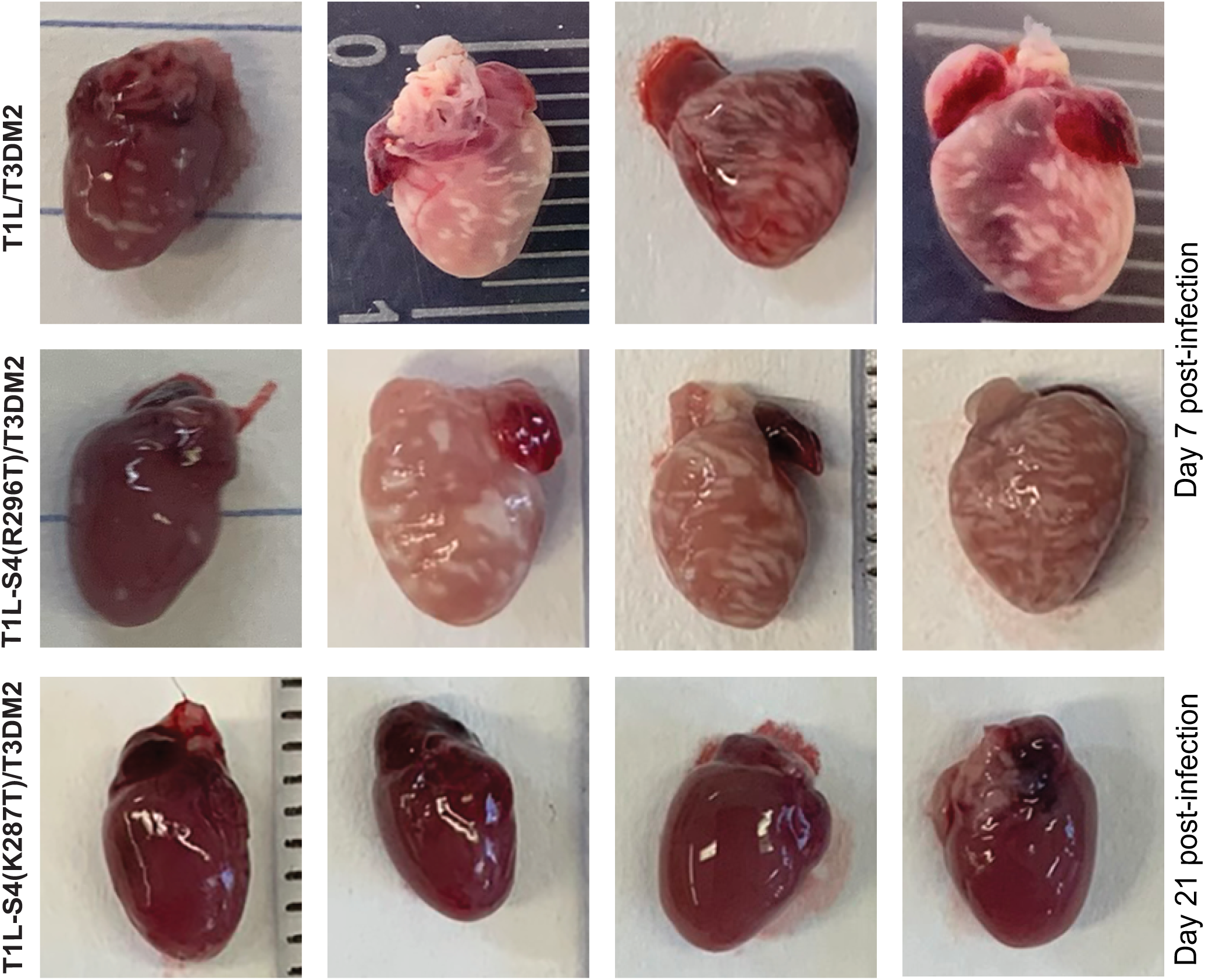
The T1L-S4(K287T)/T3DM2 reassortant causes minimal gross pathologic changes in the heart compared with the parental reassortant and T1L-S4(R296T)/T3DM2. Four representative images show the gross cardiac pathology observed in neonatal mice infected with T1L/T3DM2, T1L-S4(R296T)/T3DM2 at day 7 post-infection, and T1L-S4(K287T)/T3DM2 at day 21 post-infection.

To further characterize and quantify myocarditis, H&E-stained myocardial tissue sections from mice inoculated with T1L/T3DM2, T1L-S4(R296T)/T3DM2, and T1L-S4(K287T)/T3DM2 were examined by a board-certified veterinary pathologist on days 1, 2, 4, and 7 post-infection. Based on the level of inflammation, histological changes were scored as normal, mild, moderate, or severe (**Fig. 5 and Table 1**). At day 1 post-inoculation, the hearts appeared histologically normal across all three viruses **(Fig. 5A)**. By day 2, mild histological changes were observed in heart sections from 2 of 7 and 1 of 8 mice inoculated with T1L-S4(K287T)/T3DM2 and T1L-S4(R296T)/T3DM2, respectively **(Fig. 5B)**; no changes were observed at day 2 in the T1L/T3DM2 hearts. Similarly, at day 4, mild to moderate inflammation was observed in the hearts of at least 50% of the mice **(Fig. 5C)**. No significant differences in heart histopathology were found between groups at days 1, 2, and 4 post-inoculation. However, at day 7, mice infected with T1L-S4(K287T)/T3DM2 had significantly lower histopathological scores than those infected with T1L/T3DM2 (p < 0.0005) and T1L-S4(R296T)/T3DM2 (p < 0.0005). These results suggest that the T1L-S4(K287T)/T3DM2 virus has significantly lower virulence than the parental T1L/T3DM2 reassortant.

**Figure 5.**
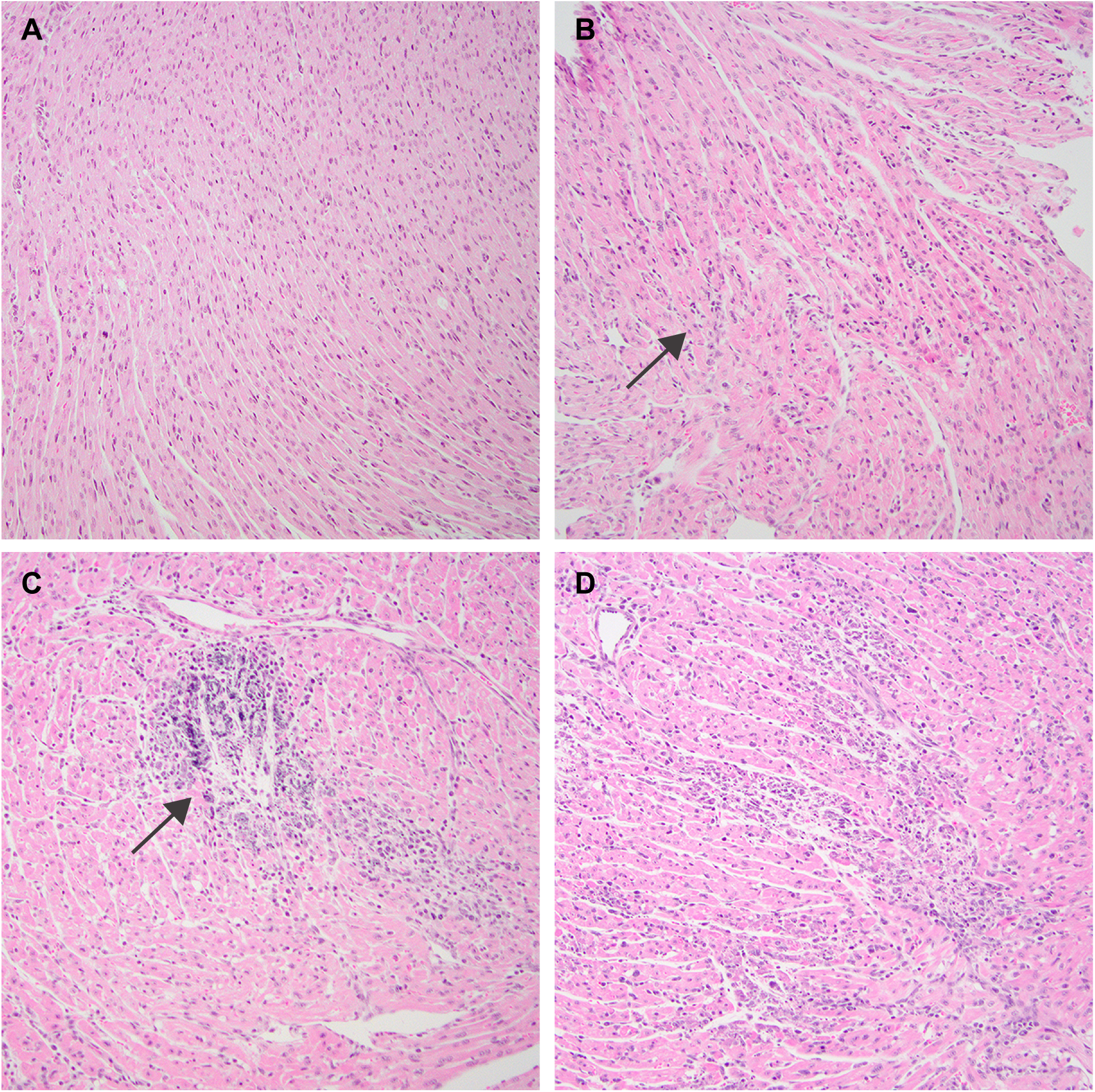
Histopathology of myocardial tissue sections from infected mice showing examples of normal, mild, moderate, and severe inflammation. Formalin-fixed hearts were collected from infected neonatal mice on Days 2, 4, and 7, embedded in paraffin, sectioned, H&E-stained, and imaged. Images were reviewed by a Board-Certified Pathologist for the presence of myocarditis (see **Table 1** for scoring). The images shown illustrate A) normal tissue,B) mild multifocal inflammation with small, discrete lesions (arrow) and some necrosis, C) moderate multifocal inflammation with larger discrete lesions (see arrow), and D) severe inflammation with multifocal-to-coalescing regions of inflammation and/or necrosis, with occasional dystrophic mineralization.

**Table 1.** Histopathological scores of infected tissue sections. Scoring: normal, mild, moderate, or severe (see **Fig. 1** for examples). ^#^ On day 1, all samples were normal. The Fisher exact test with 95% confidence was used to compare histological scores across days for the tested viruses. ^##^ Pairwise statistically significant differences were observed only on day 7 between T1L-S4(K287T)/T3DM2 and T1L/T3DM2, and between T1L-S4(R296T)/T3DM2, p < 0.0005.

| **T1L/T3DM2** | **Day 2 (n=8)<sup>#</sup>** | **Day 4 (n=13)** | **Day 7 (n=19)<sup>##</sup>** |
| --- | --- | --- | --- |
| Normal | 8 (100%) | 6 (46.2%) | 0 |
| Mild | 0 | 4 (30.1%) | 0 |
| Moderate | 0 | 3 (23%) | 0 |
| Severe | 0 | 0 | 19 (100%) |
| **T1L-S4(K287T)/T3DM2** | **Day 2 (n=7)** | **Day 4 (n=12)** | **Day 7 (n=10)<sup>##</sup>** |
| Normal | 5 (71.4%) | 6 (50%) | 0 |
| Mild | 2 (28.6%) | 6 (50%) | 7 (70%) |
| Moderate | 0 | 0 | 3 (30%) |
| Severe | 0 | 0 | 0 |
| **T1L-S4(R296T)/T3DM2** | **Day 2 (n=8)** | **Day 4 (n=10)** | **Day 7 (n=15)<sup>##</sup>** |
| Normal | 7 (87.5%) | 3 (30%) | 0 |
| Mild | 1 (12.5%) | 5 (50%) | 1 (6.7%) |
| Moderate | 0 | 2 (20%) | 1 (6.7%) |
| Severe | 0 | 0 | 13 (86.7%) |

### Distribution of viral antigen by immunofluorescence

We detected viral titers indicative of infection on days 2, 4, and 7 in the hearts of mice infected with all three viruses (**Fig. 3**). Myocarditis typically becomes evident by day 7 post-infection in mice infected with REOV strains with myocarditic potential (8). We used immunofluorescence to examine the distribution of viral antigen in the heart at days 2, 4, and 7 post-infection (**Fig. 6**). No viral antigen was observed in samples collected on day 2 post-infection. At day 4, REOV antigen was present in patches of varying sizes in the hearts of T1L/T3DM2- and T1L-S4(R296T)/T3DM2-infected mice. In contrast, only small, indistinct spots of REOV antigen were detected in the hearts of T1L-S4 (K287T)/T3DM2-infected mice at day 4. By day 7 post-infection, the distribution of viral antigen in the hearts of mice infected with the parental T1L/T3DM2 and T1L-S4(R296T)/T3DM2 was similar, featuring large, coalescing patches of antigen often arranged in striated patterns (**Fig. 6**).

**Figure 6.**
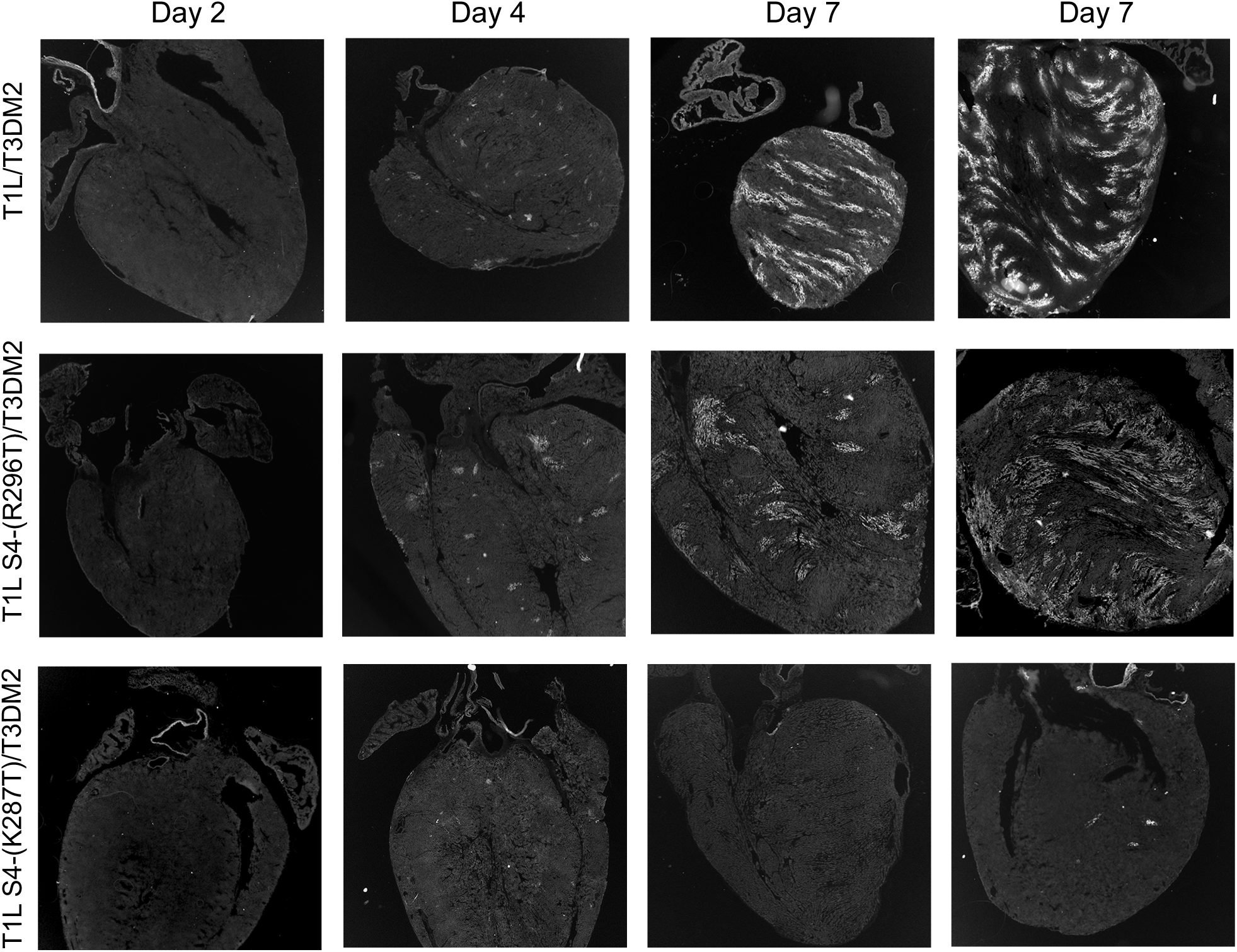
Distribution of viral antigen in infected hearts. Cryosections of heart tissue obtained from mice infected with T1L/T3DM2, T1L-S4(R296T)/T3DM2, and T1L-S4(K287T)/T3DM2 were collected at days 2, 4, and 7 pi, immunostained for REOV antigen, and imaged at 4x magnification by immunofluorescence microscopy. Representative images are shown for each time point. Contrast and brightness were adjusted to make the tissue outline visible. No viral antigen was detected at day 2 post-infection for any of the three viruses.

## DISCUSSION

Reovirus strains differ in their capacity to cause myocarditis in neonatal mice (8). The M1, M2, and S4 viral genes are known determinants of these strain differences (7). Infection with the T1L reovirus strain causes mild to moderate myocarditis, whereas T3D infection does not. However, a reassortant virus containing the M2 gene from the T3D strain in an otherwise T1L background (T1L/T3DM2) is highly myocarditic in neonatal mice, causing 90% lethality, suggesting that the μ1-encoding M2 gene is a key factor in the virus’s capacity to induce myocarditis (10). Here, we report that a single amino acid change in the σ3 protein (K287T), which prevents σ3 from suppressing PKR activation and NF-κB transcription (20, 23), significantly reduces the virulence and myocarditic potential of T1L/T3DM2. These findings indicate that the anti-innate immune functions of σ3 are crucial for virulence and pathogenesis.

After oral inoculation of neonatal mice with 1 × 10^7^ pfu of the T1L/T3DM2 recombinant virus, we observed 50% lethality (**Fig. 2**), substantially lower than the 90% lethality reported in the original study using a lower dose of 1 × 10^4^ pfu of T1L/T3DM2 (10). Consistent with this, when we used the lower inoculum dose, we replicated the 90% lethality reported by Dina Zita et al. (unpublished data). The reason for this dose-dependent difference in lethality is unknown. However, we speculate that the higher viral dose may have induced a stronger innate immune response in the intestine, limiting viral spread beyond the initial infection site and thereby reducing virulence.

The T1L-S4(K287T)/T3DM2 mutant was significantly attenuated in virulence compared with the T1L/T3DM2 parental recombinant (**Fig. 2**). The K287T mutation in σ3 is pleiotropic, affecting both its ability to inhibit PKR and its interaction with the RNA helicase (20, 23). σ3 inhibits PKR via a dsRNA-binding-independent mechanism, but the details remain unknown. In addition, WT σ3 interacts with the helicase domain of the DEAD-box/DEAH-box helicase DHX9, inhibiting its helicase activity and perturbing DHX9-dependent recruitment of RNA polymerase pol II to proximal promoters (23). These effects of σ3 on DHX9 suppress NF-κB-dependent gene transcription. Unlike WT σ3, the K287T mutant does not interact with DHX9. It seems likely that both the effects of the K287T mutation in σ3 on PKR inhibition and on its capacity to inhibit the DHX9 helicase activity contribute to the decreased virulence of the T1L-S4(K287T)/T3DM2 recombinant relative to the parental recombinant.

We previously reported that the K287T mutation in σ3 attenuated the modest cardiac damage caused by the T1L strain (4, 20). In contrast, infection with the T1L-S4(R296T) mutant induced greater cardiac damage, as assessed by Alizarin Red staining, than the WT virus, although the difference was just below the level of significance (20). Typically, all neonatal mice orally inoculated with the T1L reovirus strain survive beyond 20 days post-inoculation. In this study, we found that 33% of newborn mice orally inoculated with the T1L-S4(R296T) mutant succumbed to infection before 20 days, compared with 100% survival in mice inoculated with the T1L parent (Fig. 2). The R296T mutation in σ3 abolishes dsRNA-binding but does not prevent σ3 from inhibiting PKR (20). The reason for the increased virulence of the T1L-S4(R296T) mutant relative to T1L is unclear. The R296T mutation does not perturb σ3’s capacity to interact with DHX9 or to inhibit PKR. We have found that σ3 interacts with other host proteins essential for regulating the innate immune response (23). It is feasible that the R296T mutation may modulate σ3’s interaction with one or more of those proteins.

Following oral inoculation, the T1L strain of reovirus first enters Peyer’s patches in the ileum by binding to M (microfold) cells and gaining access via transcytosis (24, 25). From the Peyer’s patches, the virus infects intestinal epithelial cells via their basolateral surfaces, is transported to regional lymph nodes, and then spreads via lymphatics to other organs (26). Unlike the parental T1L/T3DM2 reassortant, the K287T mutant replicated to 100-fold lower levels in the ileum. Despite this significant replicative deficit in the ileum, the K287T mutant disseminated to the liver, spleen, heart, and brain, achieving titers in those organs equivalent to those of the parental strain at 2 days post-infection. Indeed, the peak titers of the K287T mutant were similar to those of the parental strain in the brain and liver, but were significantly lower than those of the parental strain in the spleen and heart by day 4 post-infection. Given that σ3 counteracts the restrictive actions of DHX9 and PKR (20, 23), this observation suggests that in the intestine, reovirus replication is constrained by DHX9 and PKR. However, the subsequent successful spread indicates that DHX9 and PKR may play a lesser role in restricting replication during spread to the brain and liver, but are important in the heart and spleen.

## ACKNOWLEDGMENTS

We thank Dr. Danica Sutherland for excellent technical assistance and advice. This research was supported by Public Health Service award R01 AI063036 (J.S.L.P.).

